# Comparative assessment of co-folding methods for molecular glue ternary structure prediction

**DOI:** 10.1101/2025.05.25.655997

**Authors:** Yiyan Liao, Jintao Zhu, Juan Xie, Luhua Lai, Jianfeng Pei

## Abstract

Molecular glues (MGs) represent an emerging therapeutic paradigm capable of inducing or stabilizing protein-protein interactions, with broad applications in creating neomorphic interactomes and targeted protein degradation. However, current discovery efforts remain largely confined to experimental screening, while *in silico* rational design of MGs persists as a formidable challenge. A critical step toward rational design lies in accurate ternary complex modeling. Given the scarcity of such complexes in the Protein Data Bank for training specialized predictive models, we tested the ability of recently developed co-folding models, including AlphaFold 3, Boltz-1, Chai-1, Protenix, and RoseTTAFold All-Atom in building the complex models. We systematically curated a dataset, named MG-PDB, with 221 non-covalent MG-engaged ternary complexes. MGBench were further introduced as a comprehensive benchmark set, which comprises 88 ternary structures excluded from co-folding models’ training data through rigorous time-based partitioning. Our benchmark results demonstrated that AlphaFold3 achieved the best overall performance among co-folding methods, in terms of both protein-protein interaction interface prediction (50.6% success rate) and MG-protein interaction recovery (32.9% success rate). However, our homology study showed that most of their successful predictions actually stemmed from memorization. Further analysis revealed three phenomena of current co-folding methods for MG ternary structure prediction. Firstly, these methods struggle to accurately model large interaction interfaces. Secondly, their predictive accuracy is notably reduced for domain-domain complexes compared to domain-motif interactions. Lastly, they face specific challenges in modeling MG degrader complexes with sufficient accuracy. We showcased they relied on the existing interaction patterns, and highlighted the need for further improved in novel E3 ligase systems. These findings reveal fundamental gaps in existing methods to learn atomic-level interaction rules for MG-engaged ternary complex modeling. MG-PDB and MGBench provide critical resources and mechanistic insights to advance computational MG discovery.

**Table of Contents Graphic:** 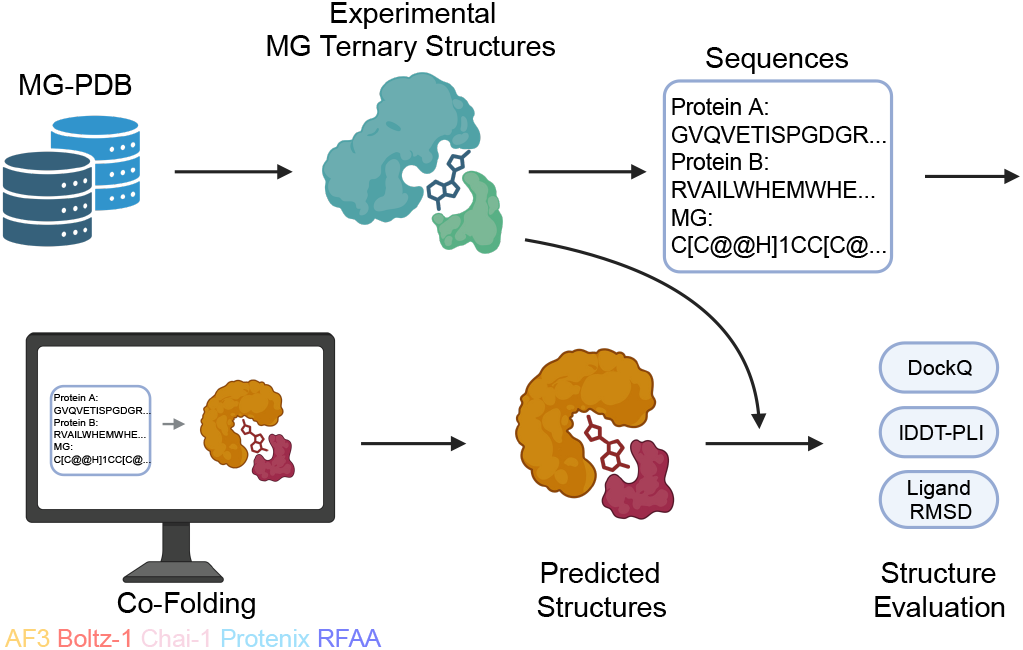

## Introduction

Chemically induced proximity has emerged as an attractive strategy in modern drug discovery, with broad biological applications in modulating protein-protein interactions (PPIs), including creating neomorphic PPIs^1^ and targeted protein degradation (TPD)^2–4^. As a central paradigm of such strategy, molecular glues (MGs) have garnered significant attention in degrader development due to their favorable drug-like properties^5–7^ and clinical success (at least nine FDA-approved drugs function as MGs)^8^. Notably, MGs provide access to challenging targets such as transcription factors^9,10^ and intrinsically disordered proteins^11^ that are traditionally considered undruggable. Mechanistically, MGs typically refer to monovalent small molecules^12,13^(with the exception of a few natural products) that either induce neomorphic PPIs or stabilize native PPIs by cooperatively binding at a PPI interface^14–16^.

Based on the mechanism of action (MOA) and downstream biological effects, MGs can be classified into three categories: molecular glue degraders (MGDs), non-degradative heterodimerizing MGs, and homodimerizing MGs^12^. MGDs represent a special class of MGs that induce the binding of target proteins to E3 ubiquitin ligases, thereby triggering ubiquitination and subsequent proteasomal degradation of the target proteins^14^. Non-degradative heterodimerizing MGs, on the other hand, modulate protein function without degradation by inducing or enhancing native PPIs^7^. Homodimerizing MGs (self-association MGs) promote protein self-association, leading to the formation of higher-order assembly, which may alter the native state of proteins and influence their function^17,18^. Despite the diversity in their MOA, all MGs mediate the formation of ternary complexes with two protein partners, which serves as the molecular basis for exerting broad regulatory effects on cellular functions.

In the past decade, MG discovery has relied on serendipity, phenotypic screening and drug repurposing^19^. Traditional methods for discovering MGs can be primarily divided into activity-based and interaction-based approaches^8^. Activity-based methods (ABMs) evaluate the effects of small molecules by measuring downstream activities in cells or *in vitro*, while interaction-based methods (IDMs) directly measure the interactions between two proteins upon the addition of small molecules. These methods typically require large-scale, high-cost experimental platforms. To accelerate the discovery of MGs and reduce costs, computational models for screening and designing of MGs need to be developed.

With the advancement of computer-aided drug design, structure-based drug design could be considered for MG discovery, enabling the exploration of a broader chemical space for effective MGs or neosubstrates^20^. However, its reliability remains a significant challenge. The structural modeling of MG-engaged ternary complexes is of significant importance for structure-based MG screening and design. Notably, any binary combination of the ternary components typically exhibits low or undetectable binding affinity, and before engagement well-defined binding pockets are not a prerequisite for complex assembly^21^. Such a cooperative binding mechanism with concurrent pocket formation necessitates modeling approaches that accommodate extensive conformational rearrangements in MG-engaged ternary complexes, exceeding the current capabilities of conventional induced-fit docking^22^ and AI-based flexible docking methods^23,24^. Recently, AI technology has achieved groundbreaking progress in biomolecular structure prediction. AI co-folding methods^25–29^ have been introduced to learn general molecular interactions and predict various biomolecular complex structures. RoseTTAFold All-Atom (RFAA)^25^ was the first to achieve a unified representation of multiple biomolecular types; AlphaFold 3 (AF3)^26^ introduced a diffusion module on the basis of its previous version AF2^30^, enabling atomic-level structure prediction. Subsequent models, including Chai-1^27^, Boltz-1^28^ and Protenix^29^, have replicated and improved upon AF3, achieving similar structure prediction capabilities. These co-folding methods perform *de novo* structure modeling from protein sequence and molecular format in SMILES, rendering them particularly suited for *in silico* determination of MG-engaged ternary complexes.

Co-folding methods have been benchmarked in several applications, including protein-ligand co-folding^31^, protein-protein complexes^32^, G protein-coupled receptor-ligand systems^33^, and PROTAC-mediated ternary complexes^34^. While demonstrating considerable modeling accuracy, these methods still rely heavily on memorization of training data^31^ and often exhibit significant errors, limiting their capability for high-precision interaction prediction. Their application to MG ternary structure modeling remains unexplored, presenting both a critical test of model generalizability and an opportunity to guide future methodological development in this emerging therapeutic paradigm.

Here, we carefully establish a dataset consisting of 221 non-covalent MG-engaged ternary complexes, named MG-PDB. Due to the lack of relevant benchmark sets, we further introduce MGBench from MG-PDB for comprehensive evaluation of MG-engaged ternary structure prediction, which includes 88 high-quality structures released after co-folding methods training. We then present a comparative assessment of five state-of-the-art (SOTA) co-folding methods (AF3, Boltz-1, Chai-1, Protenix, and RFAA) across MGBench. Our benchmark analysis reveals that AF3 outperforms other co-folding methods, achieving 50.6% prediction accuracy for protein-protein interaction interfaces and 32.9% recovery rate for MG-protein interactions. However, we observe that the memorization effect contributes significantly to this performance. Through systematic evaluation of structural determinants and mechanistic case studies, we identify critical limitations in modeling large interacting interfaces, domain-domain complexes, and degradation-specific glue mechanisms. These findings establish essential quality baselines while highlighting fundamental gaps in current approaches’ ability to capture ternary-molecule interactions. Our work provides comprehensive data resources, quantitative performance metrics and mechanistic insights to guide future development of structure prediction tools for MG discovery.

## Results

### Systematic Collection and Analysis of Experimental MG Ternary Structures

To evaluate the reconstruction capability of various co-folding methods for MG ternary complexes, we collected MG-related structures from the Protein Data Bank (PDB) database. We manually reviewed the annotated CCD IDs of MG and the protein chain identifiers directly interacting with MG, and further determined the MOAs through original literature sources. In total, we obtained 221 non-covalent MG-engaged ternary complexes, named MG-PDB, comprising 207 structures determined by X-ray diffraction and 14 structures determined by electron microscopy. A detailed analysis of MG-PDB was conducted to ensure its representativeness and high quality. Based on the classification of MOA, MG-PDB encompasses 67 MGDs, 134 non-degradative heterodimerizing MGs, and 20 homodimerizing MGs. Additionally, following the previous study^35^, 221 ternary complex structures are classified into 136 domain-domain structures and 85 domain-motif structures based on the two protein binding partners (Supplementary Table S1). Detailed statistical analysis shows the vast majority of structures have resolutions below 3.5 Å (Figure 1A), and most R-free values are below the commonly average of 0.26^36–38^ (Figure 1B), which indicates reliable structural quality. Analysis of MGs property distributions further demonstrates broad coverage of drug-like molecules within MG-PDB (Figure 1C-F), mostly conform to the characteristic features of MGs^5^. Furthermore, we assessed the number of violations to Lipinski’s rule of five, revealing that approximately 60% of MGs fully comply with the rule (Figure 1F).

**Figure 1.**
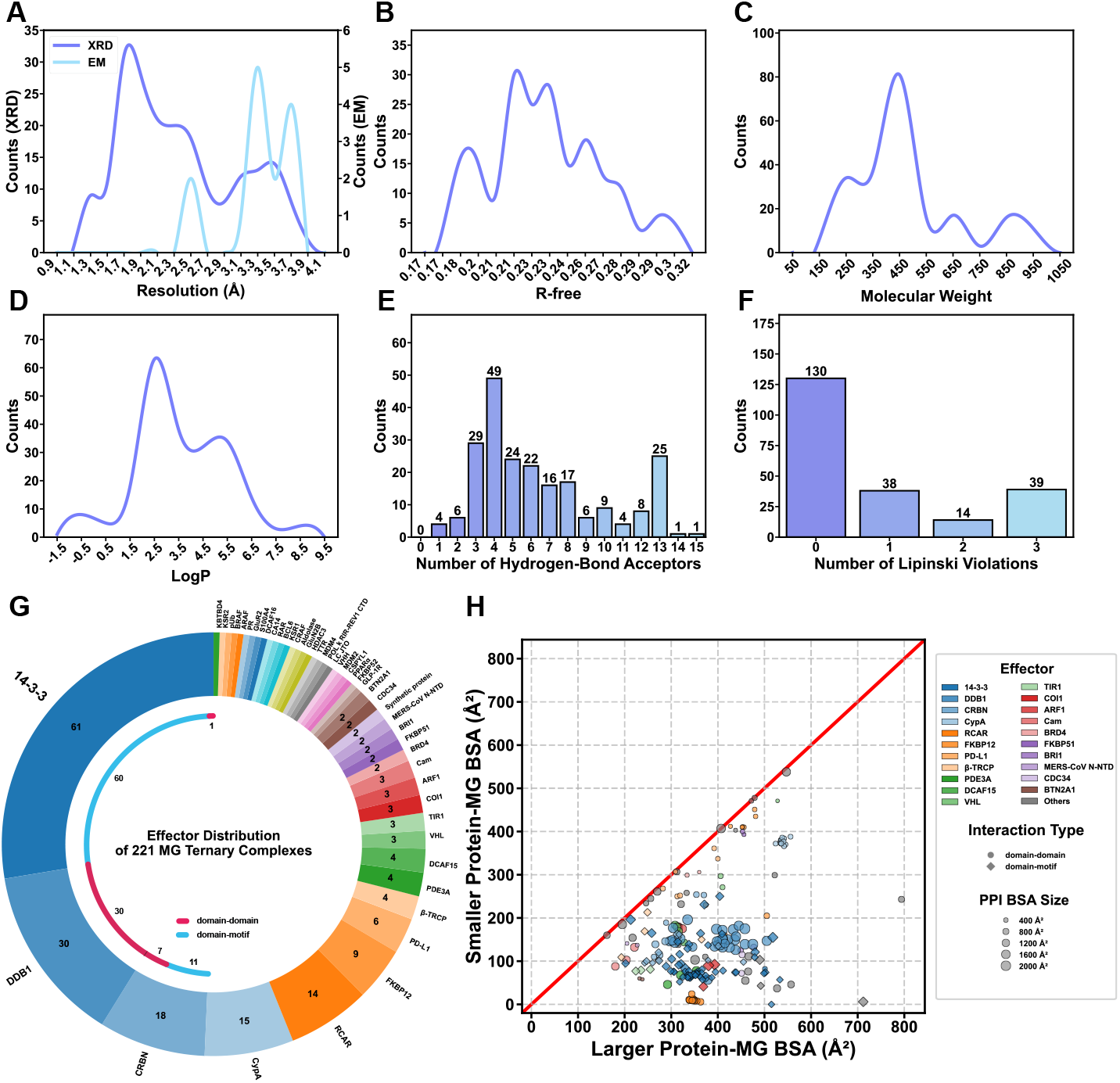
Overview of MG-PDB. For structure qualities, distribution plots showing distribution of resolution (A) and R-free value (B) for all crystal structures. For MGs molecular properties, distribution plots showing distribution of heavy molecular weight (C) and LogP (D), and histograms showing number of hydrogen-bond acceptors (E) and number of Lipinski violations (F). Doughnut chart showing the classification of the MG ternary complexes by effector protein (G). The inner circle provides a detailed breakdown of interacting modes for complexes involving 14-3-3, DDB1, or CRBN. Scatter plot showing the BSA distribution of MG ternary complexes (H). Scatters are colored by effector proteins, with the shape indicating which interacting mode the complex belong to (domain-domain: circle, domain-motif: diamond) and symbol size indicating PPI BSA of the ternary complex. XRD: X-Ray Diffraction; EM: Electron Microscopy.

A detailed analysis of the effectors involved in the ternary complexes was further conducted (Figure 1G). There are a total of 48 effector proteins in MG-PDB (excluding synthetic proteins), representing more than double the diversity reported in previous studies^35^. Among these, 14-3-3/non-degradative heterodimerizing MGs, DDB1/MGDs and CRBN/MGDs in sum account for nearly half of the whole benchmark, reflecting the current landscape of research in the MG field. Notably, most of the ternary complexes mediated by 14-3-3 or CRBN belong to the domain-motif type. Buried surface area (BSA) quantifies the extent of surface area buried upon complex formation and typically indicates the complexity of intermolecular interactions, potentially correlating with binding affinity^35,39^. We analyzed the PPI BSA as well as the two protein-MG BSAs across all ternary complexes (Figure 1H), revealing considerable diversity. The PPI BSA ranges from a minimum of 190 Å^2^ to a maximum of 2400 Å^2^, spanning a wide range. For the protein-MG BSAs, consistent with prior observations^35^, the majority of compounds exhibit asymmetric binding, where MG buries a larger surface area with one protein compared to the other. And domain-motif type complexes display a more pronounced asymmetric binding pattern as expected. Collectively, our benchmark demonstrates broad representativeness, diversity and reliability, making it well-suited for benchmarking co-folding approaches.

### A Standard Benchmark for MG Ternary Structure Modeling

To ensure the high quality of structure, we first performed a structural quality filtering on the MG-PDB dataset. Structures determined by X-ray diffraction with a resolution worse than 3.5 Å were removed. For electron microscopy structures, we further examined the local resolution representations at the MG-binding interface; if the interface resolution met 3.5 Å criteria, the structure was retained even if the overall resolution was lower than 3.5 Å. This filtering process resulted in the retention of 202 structures, this resolution-filtered MG-PDB dataset was used to perform subsequent analyses. To assess the generalization capabilities of co-folding methods, we constructed a standard benchmark utilizing structures released after 2021-09-30, called MGBench (88 structures in total). This temporal partitioning reflects evolving trends in scientific advancement and has been adopted by multiple AF3 benchmark works^31,33,34,40^. Following the interface metrics criteria for the evaluation set in AF3^26^ (as detailed in the Methods section), we performed additional homology filtering on MGBench, resulting in a final selection of 25 ternary complex structures designated as the MGBench low-homology set. The remaining 63 structures were classified as the MGBench high-homology set.

### Overall Evaluation and Homology Analysis on MGBench Reveal Memorization Effect of Current Co-folding Methods

Then, we performed ternary structure predictions using five co-folding methods, i.e., AF3, Boltz-1, Chai-1, Protenix, and RFAA on both resolution-filtered MG-PDB and MGBench. The number of successfully predicted structures for each method is presented in Supplementary Table S2. Although a few structures could not be reconstructed or cannot be evaluated due to inherent limitations of the methods, the number is small and does not impact the overall conclusions.

We conducted a comprehensive assessment of the structural quality of ternary complex reconstructions across different methods. The quality of PPI interfaces was evaluated using DockQ (Figure 2A-B and Supplementary Figure S1A-B). The DockQ distributions, except for RFAA, exhibited a similar bimodal distribution (Figure 2A and Supplementary Figure S1A), indicating that the majority of structures had either high DockQ scores (greater than 0.8) or very low scores (less than 0.1). Generally, structures with a DockQ greater than 0.23 are considered acceptable. Except for RFAA, the success rate of modeling acceptable PPI interfaces on resolution-filtered MG-PDB was approximately 70% (Figure S1B), while it dropped to around 50% for MGBench structures (Figure 2B), suggesting limited generalization capability in reconstructing PPI interfaces. The performance of RFAA was significantly inferior to other methods, and its insufficient ability to avoid intermolecular collisions was observed.

**Figure 2.**
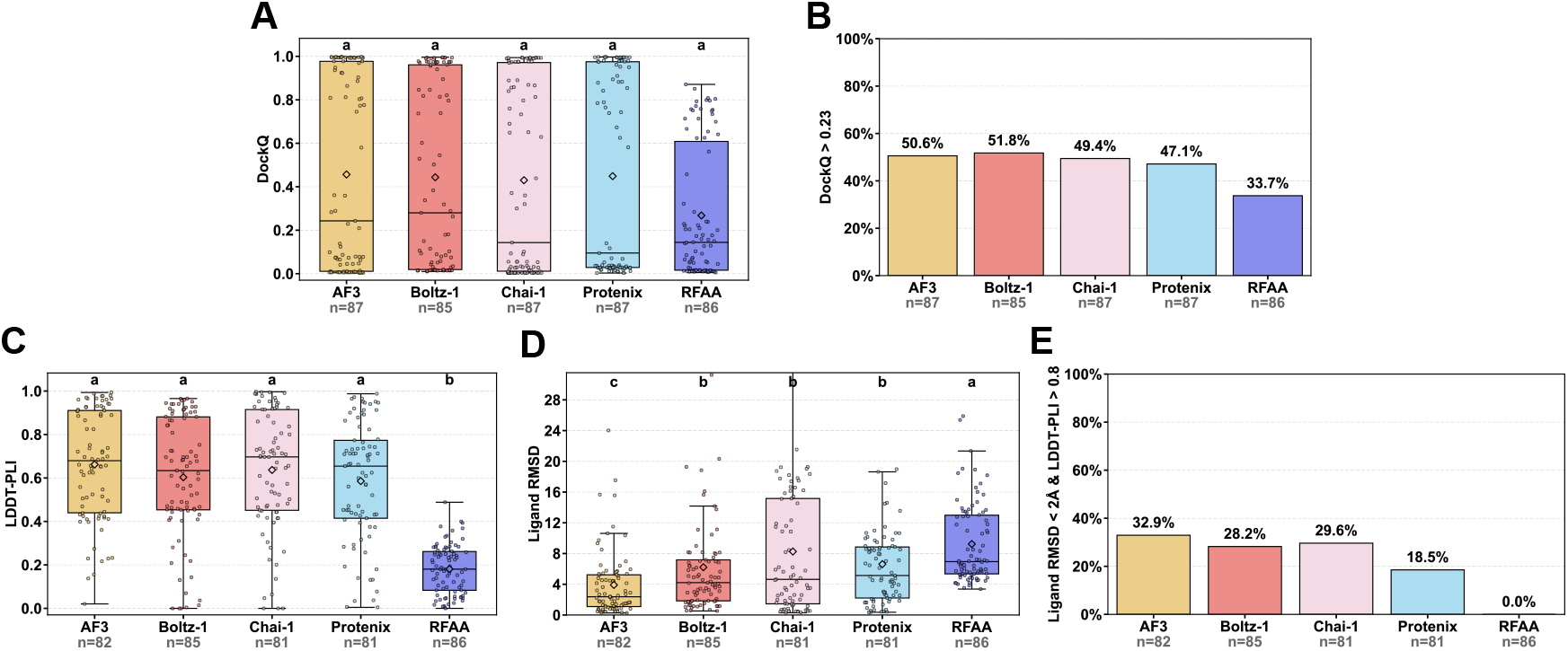
Overall metrics distributions of structural modeling in MGBench. For PPI interfaces, the box plot depicts the DockQ distribution (A) and the histogram shows the success rate defined by DockQ > 0.23 (B) for structures reconstructed by co-folding methods. For ligand-pocket binding conformations, box plots depict LDDT-PLI (C) and Ligand RMSD (D) distributions, and the histogram shows the success rate defined by ligand RMSD < 2 Å combined with LDDT-PLI > 0.8 (E) for structures generated by co-folding approaches. n represents the number of successfully modeled structures for each method. In box plots, the diamond symbol represents the mean value, and different letters indicate significant differences.

LDDT-PLI is used to evaluate atomic-level local distance deviations at protein-ligand interaction interfaces, measuring the ability of the reconstructed models to recapitulate MG-protein binding modes. Studies indicate that LDDT-PLI > 0.8 is thought to reflect a reliable interaction mode^41^. The LDDT-PLI distribution results show that for resolution-filtered MG-PDB structures, the median values of all methods (except RFAA) is around 0.8 (Supplementary Figure S1C), whereas for MGBench structures, they all fall below 0.7 (Figure 2C), indicating insufficient accuracy. Ligand RMSD assesses the precision of MG binding pose prediction. A ligand RMSD < 2 Å is generally considered indicative of native-like conformations. Results demonstrate that while the median RMSD for resolution-filtered MG-PDB structures is around this threshold (Supplementary Figure S1D), performance degrades significantly for MGBench structures (Figure 2D), indicting room for improvement to reproduce ligand binding poses. Notably, AF3 achieves significantly lower Ligand RMSD values than other methods, highlighting its superior accuracy in predicting MG binding poses.

Based on the previously established criteria for AI-based protein-ligand co-folding structure prediction^31^, we evaluated the success rates of different models in the recovery of native ligand-pocket interaction using the dual criteria of ligand RMSD < 2 Å and LDDT-PLI > 0.8. As shown in Figure 2E and Supplementary Figure S1E, the performance varied significantly among models. RFAA failed to generate any successful predictions, indicating its lower accuracy. Compared to the success rates of approximately 50% for resolution-filtered MG-PDB structures, the success rates of all models dropped to approximately 30% for MGBench structures, suggesting limited capability of these methods for native ligand-pocket interaction reconstruction. Notably, AF3 demonstrated the highest success rate (32.9%) among all models for MGBench structures, indicating its relatively stronger performance.

We further conducted a sequence homology analysis on MGBench, examining the reconstruction success rates on both MGBench high-homology set and MGBench low-homology set (Figure 3). A clear observation was that the structural reconstruction success rate on the MGBench low-homology set was significantly lower than that on the MGBench high-homology set, indicating a memorization effect across the methods. This finding aligns with conclusions from previous benchmarking^31^.

**Figure 3.**
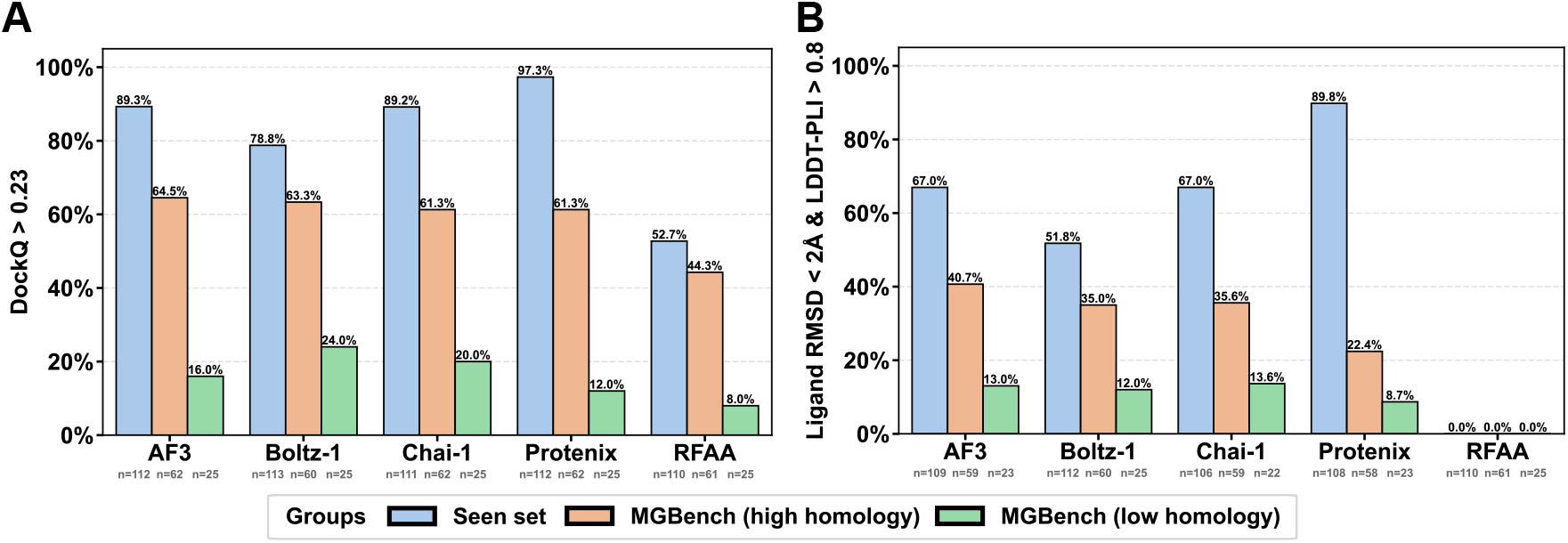
Homology analysis among co-folding methods. Histograms showing the acceptable rates of PPI interface quality (A) and recovery rate of native ligand-pocket interaction (B) on the seen set (i.e., structures in resolution-filtered MG-PDB but not in MGBench), MGBench high-homology set and MGBench low-homology set. n represents the number of successfully modeled structures for each group.

### Analysis of Key Factors Influencing MG-Engaged Ternary Complex Prediction

In the preceding section, we observed significant variability in the reconstruction performance across different ternary complex structures, suggesting that intrinsic structural properties strongly influence the outcomes. To further evaluate the impact of distinct structural features on the assessment metrics, we conducted a detailed subgroup analysis based on the following parameters: (1) the number of heavy atoms in the MG, (2) BSA of PPI interface, (3) protein interacting mode (domain-domain/domain-motif), and (4) the downstream functional type of the MG (MGDs/non-degradative heterodimerizing MGs/homodimerizing MGs).

First, to assess the model’s reconstruction capability for MGs with the heavy atom number, we analyzed success rates in the recovery of native ligand-pocket interaction as a function of heavy atom number (Figure 4A). The overall trend revealed a notable decline in prediction accuracy for MGs with larger MG molecules, indicating potential limitations in the training set’s coverage and insufficient learning of chemical space. Specifically, MGs with a large heavy atom number are often natural products, whose structural diversity and conformational complexity may introduce prediction challenges^42,43^.

**Figure 4.**
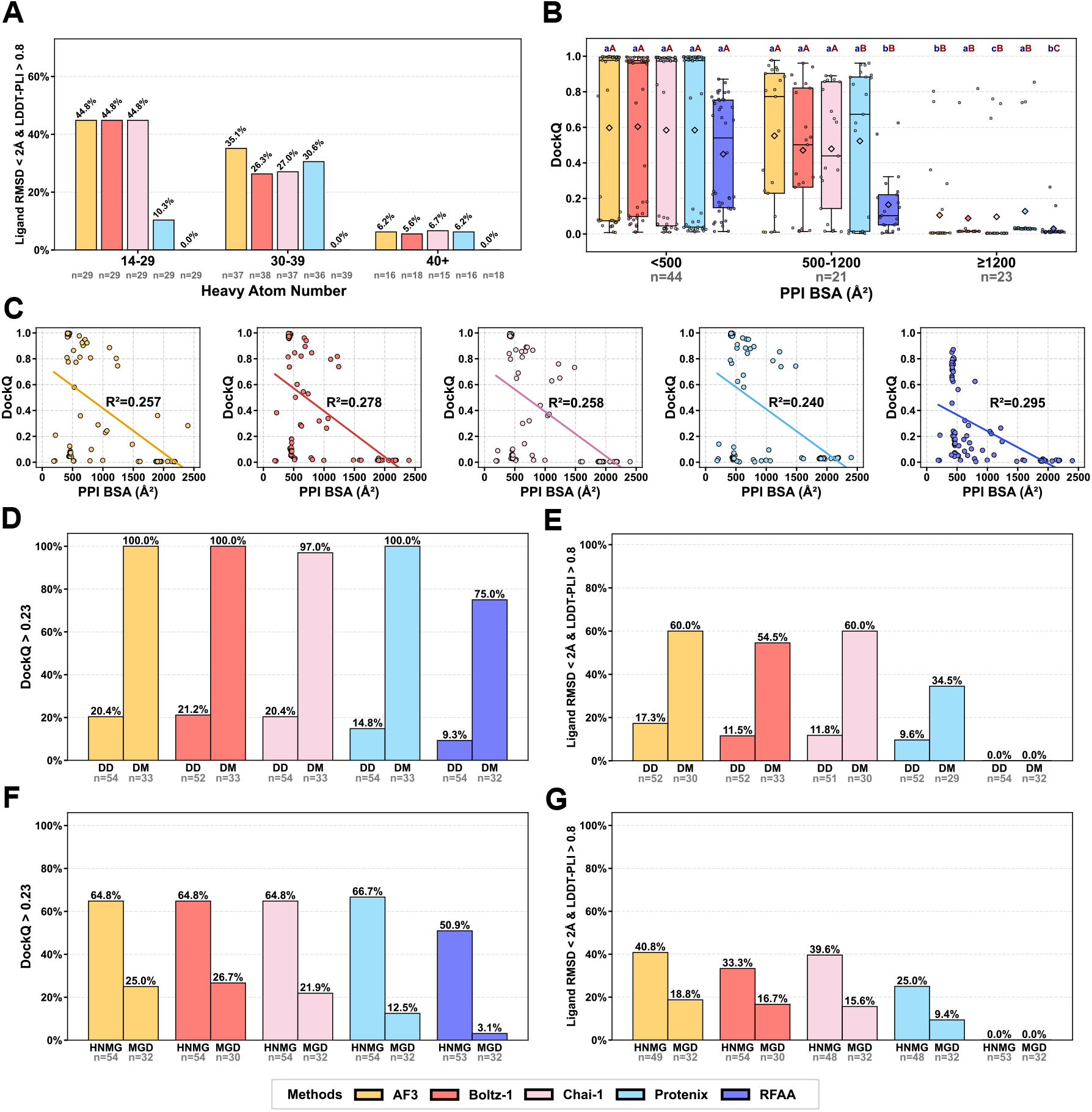
Analysis of key factors influencing MG-engaged ternary complex prediction. Histogram showing the success rate across heavy atom number for reconstructed structures from MGBench (A). Box plot showing the distribution of DockQ categorized based on BSA of native ternary complex structures (<500 Å^2^, 500-1200 Å^2^, >1200 Å^2^) (B), and scatter plots displaying the relationship between DockQ and BSA of native structures, along with the fitted regression line (C). Histograms showing the impact of interacting mode (DD: domain-domain, DM: domain-motif) on the PPI interface quality (D) and recovery of native ligand-pocket interaction (E). Histograms showing the impact of MG type (HNMG: non-degradative heterodimerizing MGs, MGD: MG degraders) on the PPI interface quality (F) and recovery of native ligand-pocket interaction (G). In the box plot, the diamond symbols represent the mean value, with lowercase letters indicating significance between different methods (same BSA group) and uppercase letters indicating significance between different BSA groups (same method). n represents the number of successfully modeled structures for each group.

The BSA of PPI interfaces typically reflects the size of the PPI interface. Previous studies have attempted to explore the correlation between BSA and binding affinity data in MG ternary complexes^35^. Here, we calculated the BSA of native MG ternary complexes and analyzed its relationship with the DockQ metric (Figure 4B-C). Within MGBench, a clear negative correlation between BSA and DockQ was observed (Figure 4B), indicating that current methods struggle to accurately model large PPI interfaces. Linear regression analysis between DockQ and BSA further confirmed this negative correlation, with all methods showing correlation coefficients of more than 0.25 (Figure 4C), reinforcing our conclusion. In summary, current co-folding methods fail to adequately model the PPI mechanisms in MG-engaged ternary complexes, and their modeling quality deteriorates significantly as the PPI interface size increases.

We investigated the impact of protein interacting mode categories on evaluation metrics. The results (Figure 4D-E and Supplementary Figure S2) demonstrate significant differences in all metrics between domain-domain and domain-motif groups across all methods, indicating that the reconstruction quality of domain-motif structures is significantly higher than that of domain-domain structures. Notably, the success rate (DockQ > 0.23) for MGBench reconstructed structures approaches 100% in all methods (except RFAA) (Figure 4D), which aligns with our previous conclusion since domain-motif structures typically feature smaller PPI interfaces and BSA. Furthermore, the native interaction recovery also shows significantly higher success rates for domain-motif structures (Figure 4E). The achieved performance is not surprising, as nearly all domain-motif structures in MGBench exhibit high homology to those in the training set, where only CRBN and 14-3-3 hub protein-mediated ternary complex systems were included (Supplementary Table S3).

To support the design of MGs with different MOAs, we separately analyzed the distribution of reconstruction metrics. The results show that the quality of reconstructed ternary complexes induced by MGDs is significantly poorer across multiple metrics (Figure 4F-G and Supplementary Figure S3), achieving only ~20% DockQ success rate and ~15% native interaction recovery rate on MGBench, respectively (Figure 4F-G). Currently, research and design of MGs primarily focus on MGDs utilizing TPD mechanisms. Here, we emphasize that current co-folding methods fail to meet the precision requirements for modeling MGD-engaged structures, making it challenging to elucidate the mechanisms of MGDs and *in silico* design.

### Case Studies for Structural Modeling of Ternary complexes between MGD-Engaged E3 Ligase and Neosubstrate

We selected representative structures from the MGBench low-homology set for detailed analysis. The WIZ zinc finger protein associates with the cohesin/CTCF complex at DNA loops involved in regulating gene expression and genome architecture^44,45^, and has been linked to the pathogenesis of sickle cell disease^10^. Recently, Novartis Biomedical Research reported the development of dWIZ-1, a thalidomide analog MGD that reprograms CRBN by inducing a CRBN/MGD neosurface for WIZ engagement to trigger its ubiquitination and degradation by the proteasome^10^. The corresponding complex structure has been solved (PDB: 8TZX). We observed that the current co-folding methods are capable of accurately modeling both the PPI interface and the pocket-ligand interactions within this complex (Figure 5). It is predictable as WIZ belongs to the Cys2-His2 (C2H2) subfamily, some of which are well-investigated zinc finger substrates targeted by CRBN-MGD complexes^46^. Although WIZ exhibits limited primary sequence similarity (<40% identity) to known thalidomide-analogs targeting C2H2 zinc fingers in our homology analysis, the structural conservation of the characteristic G-loop motif among C2H2-type zinc fingers enables specific recognition and binding by CRBN-MGD complexes^47^. Ternary structures of CRBN with different C2H2-type neosubstrate-small molecule pairs in the training set, such as CK1a-lenalidomide (5FQD), GSPT1-CC885 (5HXB), ZNF692-pomalidomide (6H0G), SALL4-thalidomide (6UML) and IKZF2-ALV1 (7LPS), can promote co-folding methods to capture the inherent binding pattern mediated by the structural conserved degron motif. As a motif alignment analysis from Monte Rosa Therapeutics predicts that there are 2,550 putative G-loop-containing proteins in the human proteome^47,48^, the successful case demonstrates that current co-folding methods could be a promising tool for accurate modeling of G-loop-containing neosubstrates engaged with CRBN-thalidomide-analog complexes.

**Figure 5.**
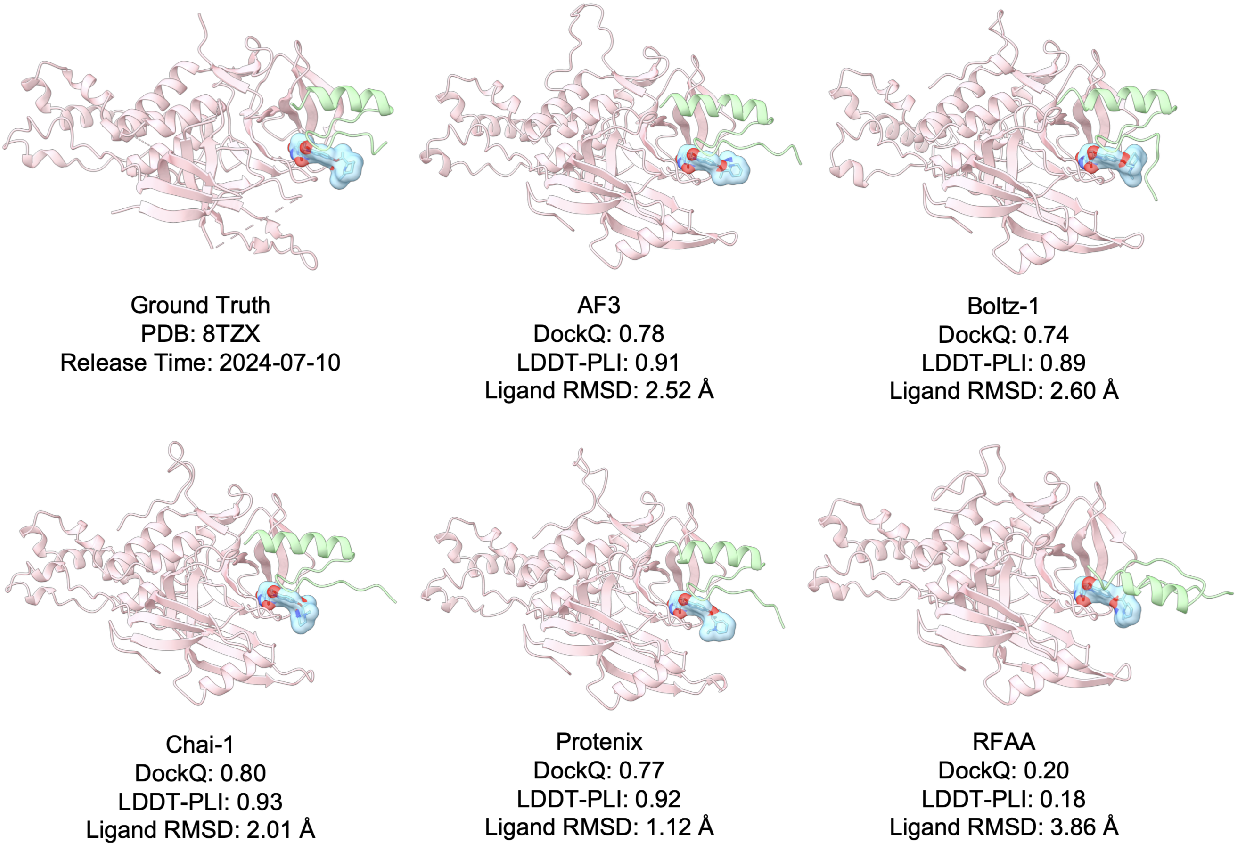
Structural modeling of the CRBN:dWIZ-1:WIZ(ZF7) ternary complex (PDB: 8TZX). The original crystal structure (top left) and computational models generated by five different co-folding methods are shown with their corresponding evaluation metrics. Proteins are shown in cartoon representation (WIZ(ZF7) in light green, CRBN in light pink), with the MGD dWIZ-1 depicted as blue sticks with transparent surface.

Given that the preceding case is limited to CRBN-mediated TPD, we sought to explore whether these methods could successfully model ternary complexes involving a broader range of E3 ligases and substrates. UM171 functions as MG, inducing high-affinity interactions between KBTBD4, a substrate receptor of the CUL3–RING E3 ubiquitin ligase complex, and histone deacetylase HDAC1/2, thereby promoting the degradation of the LSD1–CoREST corepressor complex and consequently enhancing *ex vivo* human hematopoietic stem cell self-renewal^49^. We performed a reconstruction of the KBTBD4:UM171:HDAC1 ternary complex (PDB: 8VOJ), a structure included in MGBench low-homology set. We observed that all methods exhibited poor performance in predicting the PPI interface, protein-ligand interactions, and MG conformation (Figure 6). Notably, the MG predicted by Protenix was not located within the PPI interface, and the structure predicted by RFAA displayed geometric overlapping chains. Furthermore, among the remaining four cases with novel E3 ligases, including DCAF16 and VHL, almost none of the co-folding approaches yielded high-quality ternary complex structures (Supplementary Table S4). To enhance co-folding performance for other E3 ligases, substantial improvements are still needed in future work.

**Figure 6.**
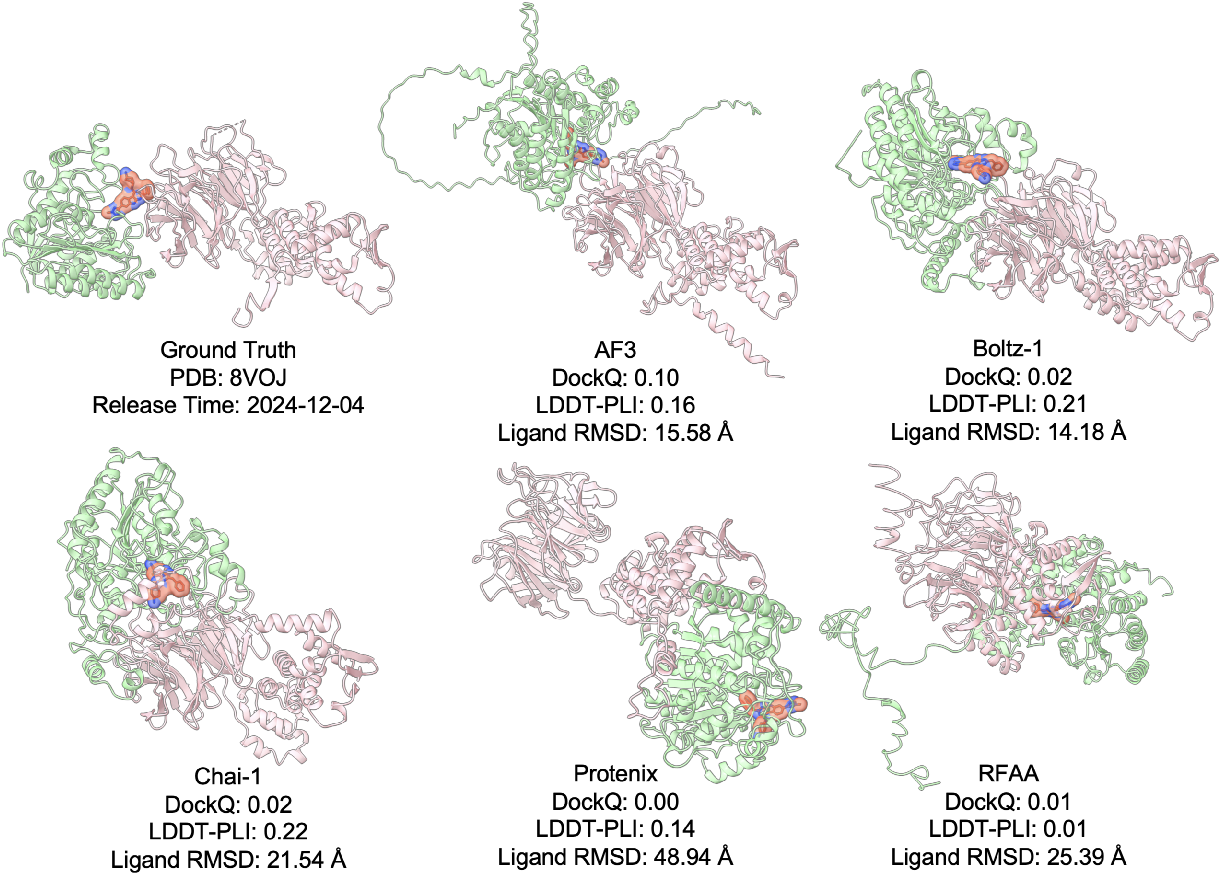
Structural modeling of the KBTBD4:UM171:HDAC1 ternary complex (PDB: 8VOJ). The original crystal structure (top left) and computational models generated by five different co-folding methods are shown with their corresponding evaluation metrics. Proteins are shown in cartoon representation (HDAC1 in light green, KBTBD4 in light pink), with the MGD UM171 depicted as red sticks with transparent surface.

## Discussion

MGs have garnered significant attention due to their favorable druggability and unique MOA^50^. Traditional discovery approaches primarily rely on serendipitous findings and large-scale experimental screening. The structural basis provided by molecular glue ternary complexes is pivotal for mechanistic studies, structure-based virtual screening, *de novo* design, and lead optimization. Emerging AI co-folding methods^25–29^ enable *de novo* prediction of these complexes from sequence and chemical data, overcoming limitations of conventional approaches^22–24^ in modeling MG-induced conformational changes. However, the modeling capabilities of these methods on MG-engaged ternary complexes remains unclear. Our study systematically assesses the performance and generalization ability of co-folding methods for a possible shift to the next-generation MG discovery paradigm.

Previous collections of MG-engaged ternary complex structures were limited to fewer than 100 entries^35^, which is insufficient for comprehensive evaluation, and most of them were used to train current co-folding methods. We systematically collected a dataset containing 221 experimentally determined non-covalent MG-engaged ternary complex structures, named MG-PDB, and conducted benchmarking study over five co-folding methods in our newly introduced benchmark set, named MGBench. Multiple quality assessment metrics were calculated, including DockQ, LDDT-PLI, and Ligand RMSD. Overall, all methods demonstrated limited reconstruction capability for MGBench structures, with approximately 50% success rate for PPI interfaces and ~30% native interaction recovery rate. Specifically, RFAA failed to predict any structures successfully, whereas AF3-like methods consistently generated reliable models. AF3 exhibited superior performance across all evaluation metrics, demonstrating its SOTA performance. Further key impact analysis showed performance decoy was observed in larger MGs, gluable PPI with large BSA and domain-domain-type PPI. The benchmark results revealed current methods remain unable to generalize predictions for novel ternary complexes, failing to adequately capture intricate molecular interaction patterns. These findings aligned with recent AF3 benchmark study in protein-ligand co-folding^31^, collectively highlighting fundamental gaps and memorization issue in accurate modeling of protein-ligand complex structures.

The primary obstacle preventing current co-folding methods from accurately predicting MG-bound ternary complex structures stems from severe data scarcity. The PDB contains dramatically fewer “protein-small molecule-protein” interaction entries compared to abundant PPI data^51^. Consequently, these methods, trained on minimal MG-PPI examples, inevitably fail. Furthermore, their AlphaFold-like architectures heavily rely on co-evolutionary information encoded in multiple sequence alignments to guide PPI modeling^26,30,52^, while some MG-PPIs lack such intrinsic evolutionary constraints since MGs typically induce neomorphic PPIs. Notably, existing successful MG design cases predominantly target systems with inherent co-evolutionary relationships, primarily thalidomide-analogs MGDs targeting CRBN-G-loop degrons^47,53^ and 14-3-3/client protein stabilizers^54^, where weak endogenous PPIs and biological function upon binding pre-exist^21^. This aligns with our benchmark results showing superior performance on domain-motif-type PPIs and certain MGD targets, as these systems retain evolutionarily coupled interaction signatures.

Beyond high-accuracy ternary complex modeling, developing scoring functions to evaluate binding affinity or degradation efficiency is urgent for computational MG screening. While AI-based scoring function remains impractical due to data scarcity, hybrid physical/AI approaches, such as GlueMap^55^ and MOLDE^56^, which integrate molecular dynamics simulations, binding free energy calculations, and generative AI, have shown promising performance in retrospective validations. An alternative strategy leverages interaction data to train classifiers, exemplified by MaSIF-neosurf’s innovative use of biomolecular surface patches to predict binding propensity in ternary systems^57^. However, this approach requires a *priori* strong affinity and stable binding poses between the MG and at least one protein partner, effectively reducing the ternary problem to binary interaction prediction between protein-ligand neosurfaces and protein surfaces, a simplification that limits generalizability. Notably, the confidence scores output by AF3-like models warrant investigation as potential affinity proxies, representing an attractive avenue for future exploration given their intrinsic structural insights.

Despite these challenges, a divide-and-conquer approach leveraging AF3 remains viable. For CRBN-G-loop systems, AF3 reliably generates structural models that can accelerate the discovery of novel MGDs. However, targeting alternative E3 ligases or developing MGDs with distinct MOAs will require systematic accumulation of experimental data, and deeper mechanistic studies. A critical open question is whether other E3 ligases exhibit conserved degron-recognition motifs analogous to the CRBN-G-loop interaction pattern. Through iterative integration of experimental data and conserved interaction constraints, continuously refined AF3 models will emerge as a transformative tool for both MG discovery and mechanistic elucidation.

Overall, while AI biomolecular co-folding methods have advanced MG-engaged ternary complex structure prediction, our benchmark study reveals their limitations and provides important references for future method optimization and rational MG design, thereby advancing computation-driven next-generation MG discovery paradigms.

## Methods

### Data Collection

We conducted a systematic search of complex structures on PDB using MG-related keywords, collecting and annotating all the structures^38^. Specifically, we manually downloaded the mmcif files and fetched structural related information through RCSB PDB Data API. To facilitate analysis of protein pairs directly interacting with MGs, we systematically annotated the two protein chain identifiers that establish direct interactions with the MG in each PDB entry, as well as the CCD code of MGs. All entries were rigorously validated as authentic MG complexes and annotated with MOA class through original literature, ensuring dataset reliability for downstream benchmarking. Notably, this study exclusively focuses on non-covalent MG-induced ternary complexes. For each PDB entry, we programmatically retrieved directly interacting protein partners’ sequences through the RCSB PDB Data API, while manually curating canonical SMILES representations of MGs - both serving as foundational input for structural reconstruction workflows. Following the category rule established by Rui et al.^35^, we further categorized and annotated the collected complex structures into two distinct domain types: (1) domain–domain: characterized by interactions between well-structured protein domains; (2) domain-motif: defined by one of the binding partners being a short linear motif containing a specific recognition sequence.

### MGBench Curation

For the 221 ternary complex PDB structures we collected, we first removed structures determined by X-ray diffraction (XRD) with a resolution worse than 3.5 Å. For electron microscopy (EM) structures, we further examined the local resolution representations at the MG-binding interface; if the interface resolution met 3.5 Å criteria, the structure was retained even if the overall resolution was lower than 3.5 Å. We further retained structures with a release date after 2021-09-30 as MGBench including 88 structures, excluding structures used in the training sets of the methods. To further investigate the impact of sequence homology between MGBench and co-folding methods’ training data, we extracted sequences from all PDB structures released before 2021-09-30 as training data, and performed a similarity search using MMSeqs2^58^ on all sequences in MGBench. Following AF3’s approach for evaluating interface metrics^26^, we applied the following filtering criteria:

- If both polymers have a length of at least 16 amino acids and exhibit greater than 40% sequence identity to two chains in the same complex in the training data, the structure is filtered out.
- For structures containing chains with fewer than 16 residues, the similarity of the longer chain must be less than 40% sequence identity to any structure in the training data.

Finally, we retained 25 structures as the MGBench low-homology set, while the remaining 63 structures were designated as the MGBench high-homology set.

### Ternary Structure Prediction

We evaluated five SOTA co-folding methods for MG ternary complex reconstruction: AF3^26^, Chai-1^27^ (https://lab.chaidiscovery.com/), Boltz-1^28^, Protenix^29^ (https://protenix-server.com), and RFAA^25^. A summary of the different methods is provided in the Supplementary Methods. To ensure a rigorous evaluation of their maximum predictive performance, all methods were implemented under optimal configurations, including the use of multiple sequence alignments and default parameter settings. For Chai-1 and Protenix, predictions were generated via their real-time updated web servers to leverage the most current implementations. The remaining methods (AF3, Boltz-1, and RFAA) were run locally using the latest software versions. Structural visualization and comparative analysis were performed using ChimeraX-1.9^59^.

### Ternary Structure Evaluation

We systematically evaluated reconstruction accuracy through comparative analysis between predicted models and experimentally determined PDB structures. The assessment employed three established metrics: (1) DockQ score for protein-protein interface quality assessment^60^, (2) LDDT-PLI (Local Distance Difference Test for Protein-Ligand Interactions) to assess the recall of contacts between ligands and pockets, and (3) ligand RMSD to measure the accuracy of ligand binding pose prediction. Specifically, the formulas for the three metrics are defined as follows:

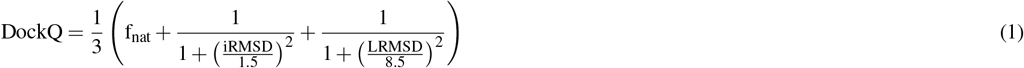

Here, f_nat_ quantifies the fraction of correctly predicted interfacial contacts, iRMSD measures structural divergence at the binding interface using backbone atoms RMSD, and LRMSD evaluates global structural alignment by computing the backbone deviation between predicted and native structures after optimal superposition.

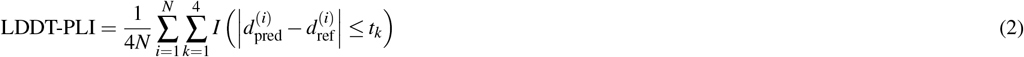

Here, *N* represents the number of effective atom pairs within the inclusion radius between ligand atoms and surrounding protein atoms in the reference structure. 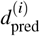denotes the distance between the *i*-th ligand-pocket atom pair in the reference structure, 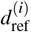is the corresponding atomic pair distance in the predicted structure, *t*_*k*_ is the preset threshold, and *I*(*θ*) is the indicator function.

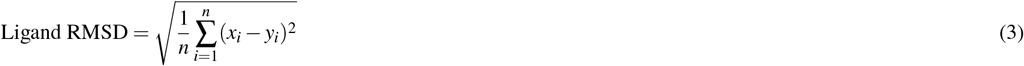

Here, *n* represents the number of heavy atoms in the ligand, *x*_*i*_ denotes the coordinate of the *i*-th atom in the ligand for the predicted conformation, and *y*_*i*_ signifies the coordinate of the *i*-th atom in the ligand for the ground truth conformation.

Computational workflows were implemented using specialized tools: DockQ calculations utilized the official DockQ package^60^, while LDDT-PLI and ligand RMSD metrics were computed through OpenStructure’s built-in validation toolkit^61^.

### Calculation of the BSA

The BSA was determined utilizing ChimeraX-1.9^59^. All solvent molecules, ligands other than MGs, and irrelevant protein chains were removed. Unique sequence identifiers were assigned to the ligands, distinct from those of the proteins. The “interfaces” command was then employed to calculate BSA for each pair of interacting chains within the complexes. We proceeded with further analysis using the BSA values obtained from PPIs.

## Acknowledgements

This work has been supported in part by the National Natural Science Foundation of China (22033001 and 32270689), the National Key R&D Program of China (2023YFF1205103), the Chinese Academy of Medical Sciences (2021-I2M-5-014) and the Anhui’s Plans for Major Provincial Science&Technology Projects (202303a07020009). We thank the Computing Platform of the Center for Life Science (Peking University) for providing resources for the GPU-based model inference. Part of the computation was performed on the computing platform of the Infinite Intelligence Pharma Ltd.

## Author Contributions

J.Z. designed the research. Y.L. and J.Z. collected the benchmark set. Y.L. conducted the experiments and analyzed the data. Y.L., J.Z. and J.X. discussed the results. Y.L. and J.Z. wrote the manuscript. J.P., L.L. and J.Z. supervised the project. J.P., L.L. and J.X. revised the manuscript. All authors read and approved the final manuscript.

## Code and Data Availability

Code for reproducing the analysis and plots presented in this paper is available at https://github.com/yiyanliao/MGBench. The data of MG-PDB and MGBench will be publicly available as soon as possible after our paper has been published.

## Supplementary Methods

### Co-folding Methods

Co-folding methods constitute a transformative advance in predicting 3D structures of molecular complexes directly from sequence data, providing unprecedented insights into biomolecular interactions. This section will summarize co-folding methods evaluated in this work.

### AlphaFold 3 (AF3)

AlphaFold 3 (AF3)^26^, proposed by the DeepMind team, is designed to model the joint structure of biomolecular complexes including proteins, nucleic acids, small molecules, ions, and modified residues. Compared to AlphaFold 2 (AF2)^30^, AF3 reduces reliance on multiple-sequence alignment (MSA) processing by replacing the evoformer module with a simpler pairformer module. It also introduces a diffusion module that directly predicts raw atom coordinates. These architectural changes enable the modeling of more diverse and general chemical structures. Notably, no structural data after 2021-09-30 was used in the training of AF3. The model achieved SOTA performance across a range of tasks. However, the AF3 model is not publicly available for training, which limits opportunities for further improvements and extensions by the broader research community.

### Chai-1

Chai-1^27^ is another state-of-the-art co-folding model developed by the Chai Discovery Team, demonstrating strong performance across a wide range of benchmarks. A key distinction from AF3 is that Chai-1 trained a unified model, rather than relying on separate models for different prediction tasks. Furthermore, Chai-1 supports the use of language model embeddings in place of MSAs, and allows the incorporation of constraint features during inference. It is worth noting that Chai-1 was trained on PDB data with a cutoff date of 2021-01-12, earlier than AF3’s training data cutoff of 2021-09-30. Chai-1 is not publicly retrainable, limiting its flexibility for further customization or adaptation.

### Boltz-1

Boltz-1^28^ is a fast-follow implementation inspired by AF3, developed by the MIT Jameel Clinic. Building upon the AF3 architecture, Boltz-1 introduces several key improvements, including a more efficient and robust MSA pairing mechanism, structure cropping during training, conditioned structure prediction based on user-defined binding pockets, modifications to the representation flow and diffusion procedures, and revision of the confidence model. Boltz-1 was trained using the same PDB data cutoff as AF3 (2021-09-30) and achieves performance comparable to both AF3 and Chai-1. Notably, Boltz-1 provides open-source retraining scripts, offering greater flexibility and adaptability for future research and applications.

### Protenix

Protenix^29^, proposed by the AM Lab for Science team at ByteDance, provides a comprehensive and faithful reproduction of AF3. Through refining ambiguous implementation steps, correcting typographical errors, and making targeted adjustments, Protenix achieves strong performance in structure prediction across diverse molecular types. The model was trained using PDB data released before 2021-09-30, the same time cutoff as AF3, ensuring a fair comparison in training data availability. Notably, Protenix has fully open-sourced its model weights, inference code, and training scripts, significantly enhancing its accessibility and reproducibility for the broader scientific community.

### RoseTTAFold All-Atom (RFAA)

RFAA^25^, proposed by the Baker Lab at the University of Washington, was the first method to achieve a unified representation across multiple biomolecular types. Building upon the RoseTTAFold2 framework^62^, RFAA integrates one-dimensional (1D) sequence information, two-dimensional (2D) pairwise distance data derived from homologous templates, and three-dimensional (3D) coordinate information as input to generate accurate structural predictions. The model was trained on PDB data deposited before 2020-04-30. However, a notable limitation of RFAA is that the predicted 3D atomic positions often contain steric conflicts and clashes, reducing its overall efficiency and reliability in practical applications.

## Supplementary Figures

**Figure S1.**
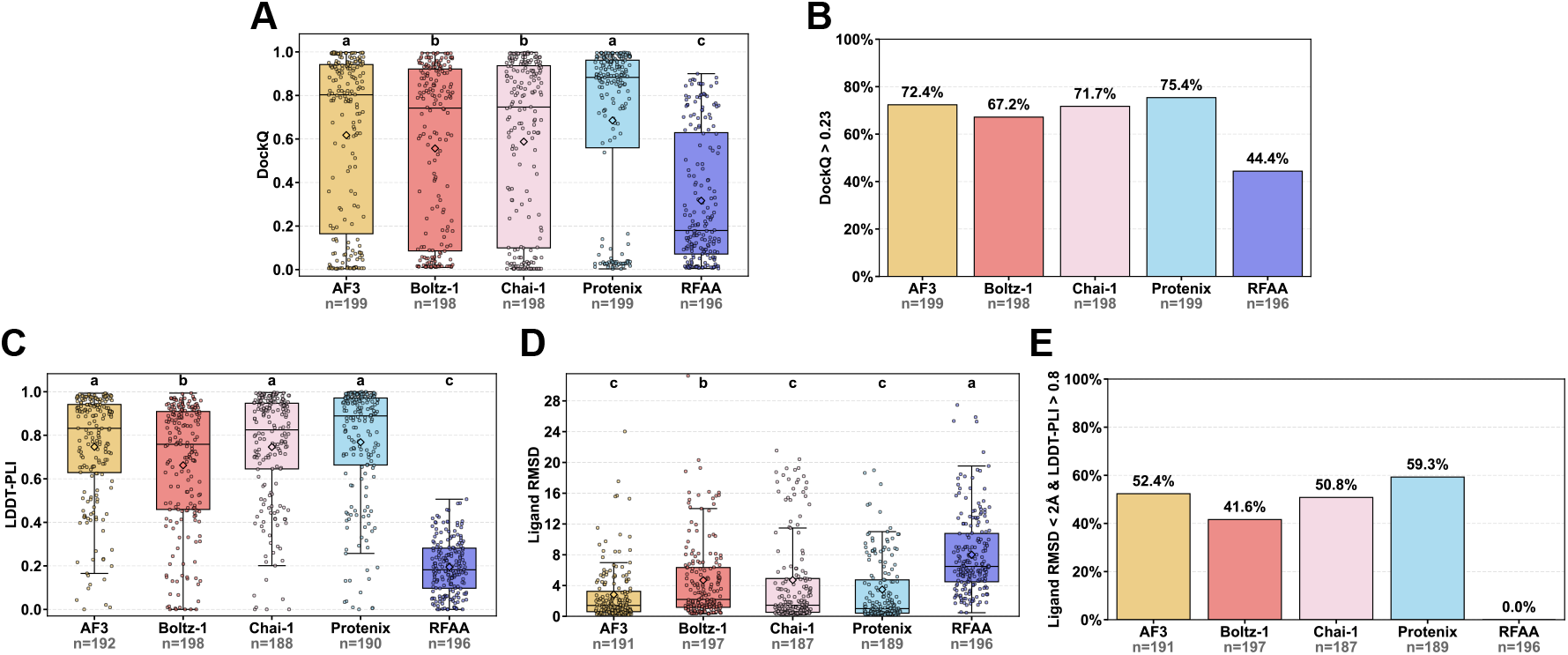
Overall metrics distributions of structural modeling for resolution-filtered MG-PDB dataset. For PPI interfaces, the box plot depicts the DockQ distribution (A) and the histogram shows the success rate defined by DockQ > 0.23 (B) for structures reconstructed by co-folding methods. For ligand-pocket binding conformations, box plots depict LDDT-PLI (C) and Ligand RMSD (D) distributions, and the histogram shows the success rate defined by ligand RMSD < 2 Å combined with LDDT-PLI > 0.8 (E) for structures generated by co-folding approaches. n represents the number of successfully modeled structures for each method. In box plots, the diamond symbol represents the mean value, and different letters indicate significant differences.

**Figure S2.**
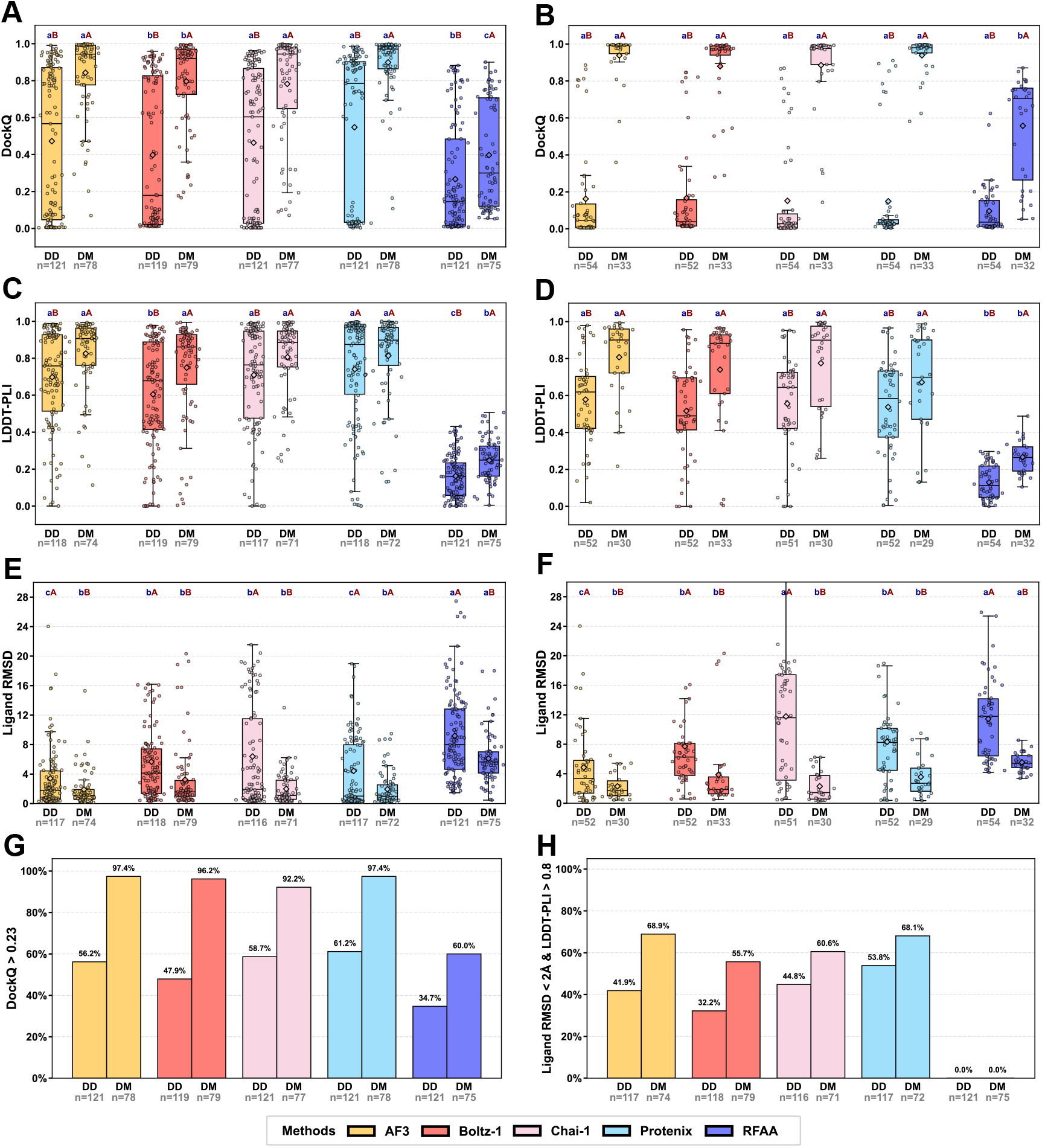
Relationship between evaluation metrics and interacting mode. Box plots showing the distribution of DockQ (A, B), LDDT-PLI (C, D) and Ligand RMSD (E, F), where left ones are evaluated in resolution-filtered MG-PDB and right ones in MGBench. Histograms showing the impact of interacting mode on the PPI interface quality (G) and recovery of native ligand-pocket interaction (H) in resolution-filtered MG-PDB structures. n represents the number of successfully modeled structures for each group. In box plots, the diamond symbol represents the mean value, with lowercase letters indicating significance between different methods (same group) and uppercase letters indicating significance between different groups (same method). DD, domain-domain; DM, domain-motif.

**Figure S3.**
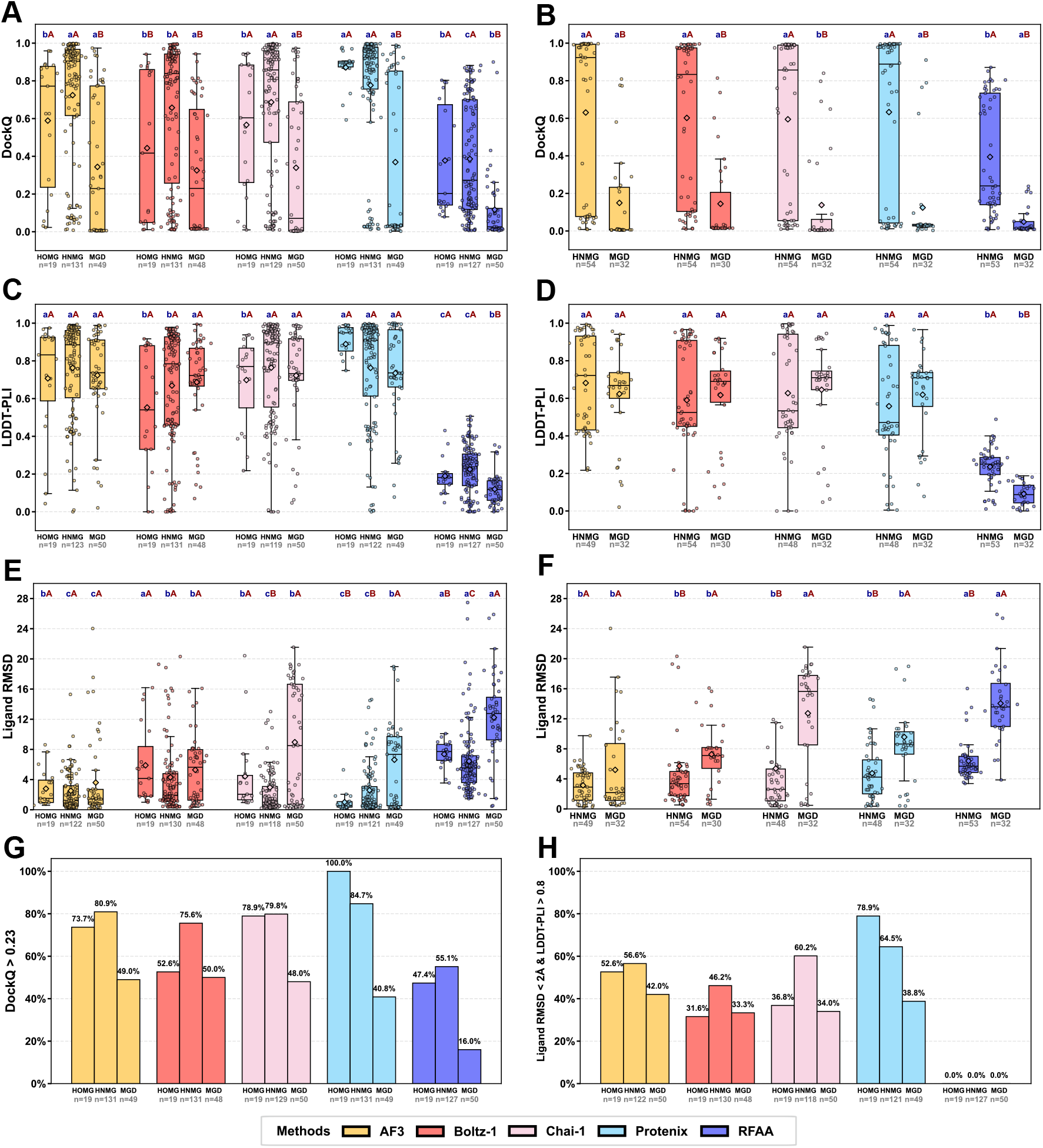
Relationship between evaluation metrics and MG type. Box plots showing the distribution of DockQ (A, B), LDDT-PLI (C, D) and Ligand RMSD (E, F), where left ones are evaluated in resolution-filtered MG-PDB and right ones in MGBench. Histograms showing the impact of MG type on the PPI interface quality (G) and recovery of native ligand-pocket interaction (H) in resolution-filtered MG-PDB structures. n represents the number of successfully modeled structures for each group. In box plots, the diamond symbol represents the mean value, with lowercase letters indicating significance between different methods (same group) and uppercase letters indicating significance between different groups (same method). HOMG, homodimerizing MGs; HNMG, non-degradative heterodimerizing MGs; MGD, MG degraders.

## Supplementary Tables

**Table S1.** Summary of dataset information.

|  | Total<br>(No.) | Class |  |  | Domain Type |  | Method |  |
| --- | --- | --- | --- | --- | --- | --- | --- | --- |
|  |  | MGD | HNMG | HOMG | DD | DM | XRD | EM |
| MG-PDB | 221 | 67 | 134 | 20 | 136 | 85 | 207 | 14 |
| MGBench | 88 | 32 | 55 | 1 | 55 | 33 | 81 | 7 |

Domain Type: DD (domain-domain), DM (domain-motif)
Method: EM (Electron Microscopy); XRD (X-ray Diffraction)
Class: HNMG (non-degradative heterodimerizing MGs); HOMG (homodimerizing MGs); MGD (MG degraders)

**Table S2.** Counts of successfully reconstructed MG ternary complexes among co-folding methods.

| Method | Resolution-filtered MG-PDB | MGBench |
| --- | --- | --- |
| AF3 | 199 | 87 |
| Boltz-1 | 198 | 85 |
| Chai-1 | 198 | 87 |
| Protenix | 199 | 87 |
| RFAA | 196 | 86 |
| Total | 202 | 88 |

**Table S3.** Summary of domain-motif structures in resolution-filtered MG-PDB & MGBench.

|  | domain-motif<br>(No.) | Effectors |  | Homology |  |
| --- | --- | --- | --- | --- | --- |
|  |  | CRBN | 14-3-3 | High | Low |
| Resolution-filtered MG-PDB | 80 | 6 | 60 | - | - |
| MGBench | 33 | 3 | 30 | 32 | 1 |

**Table S4.** Metrics of Novel E3 Ligases Ternary Complexes.

| PDB ID | E3 ligase | Method | DockQ | LDDT-PLI | Liangd RMSD |
| --- | --- | --- | --- | --- | --- |
| 8OV6 | DCAF16 | AF3 | 0.01 | 0.02 | 24.02 |
|  |  | Boltz-1 | 0.03 | 0.07 | 15.66 |
|  |  | Chai-1 | 0.05 | 0.05 | 17.39 |
|  |  | Protenix | 0.02 | 0.08 | 17.17 |
|  |  | RFAA | 0.03 | 0.02 | 25.89 |
| 8VLB | VHL | AF3 | 0.01 | 0.68 | 9.33 |
|  |  | Boltz-1 | 0.02 | 0.72 | 16.09 |
|  |  | Chai-1 | 0.01 | 0.71 | 8.46 |
|  |  | Protenix | 0.01 | 0.71 | 18.98 |
|  |  | RFAA | 0.05 | 0.03 | 19.05 |
| 8EWV | VHL | AF3 | 0.21 | 0.63 | 5.24 |
|  |  | Boltz-1 | 0.38 | 0.67 | 5.07 |
|  |  | Chai-1 | 0.09 | 0.34 | 12.32 |
|  |  | Protenix | 0.14 | 0.52 | 7.97 |
|  |  | RFAA | 0.21 | 0.12 | 8.51 |
| 8VL9 | VHL | AF3 | 0.01 | 0.62 | 11.50 |
|  |  | Boltz-1 | 0.03 | 0.65 | 13.74 |
|  |  | Chai-1 | 0.02 | 0.66 | 8.54 |
|  |  | Protenix | 0.91 | 0.97 | 0.39 |
|  |  | RFAA | 0.05 | 0.01 | 18.91 |

## Notes

### Competing Interest Statement

The authors have declared no competing interest.

